# GFF3sort: a novel tool to sort GFF3 files for tabix indexing

**DOI:** 10.1101/145938

**Authors:** Tao Zhu, Chengzhen Liang, Zhigang Meng, Sandui Guo, Rui Zhang

**Author notes:** Correspondence Rui Zhang Sandui Guo.

## Abstract

**Background:** The traditional method of visualizing gene annotation data in JBrowse is converting GFF3 files to JSON format, which is time-consuming. The latest version of JBrowse supports rendering sorted GFF3 files indexed by tabix, a novel strategy that is more convenient than the original conversion process. However, current tools available for GFF3 file sorting have some limitations and their sorting results would lead to erroneous rendering in JBrowse.

**Results:** We developed GFF3sort, a script to sort GFF3 files for tabix indexing. Specifically designed for JBrowse rendering, GFF3sort can properly deal with the order of features that have the same chromosome and start position, either by remembering their original orders or by conducting parent-child topology sorting. Based on our test datasets from seven species, GFF3sort produced accurate sorting results with acceptable efficiency compared with currently available tools.

**Conclusions:** GFF3sort is a novel tool to sort GFF3 files for tabix indexing. We anticipate that GFF3sort will be useful to help with genome annotation data processing and visualization.

## Background

As a powerful genome browser based on HTML5 and JavaScript, JBrowse has been widely used since released in 2009[1, 2]. According to its configuration document[3], it works by first converting genome annotation data in GFF3 file formats to JSON files by a built-in script “flatfile-to-json.pl”, and then rendering visualized element models such as genes, transcripts, repeat elements, etc. The main problem, however, is that this step is extremely time-consuming. The time is proportional to the number of feature elements in GFF3 files (Additional file 1). Even for small genomes like yeast (*Saccharomyces cerevisiae*), it takes ∼10 seconds to finish the conversion. For large and deeply annotated genomes such as that of humans, the time increases to more than 15 minutes. In addition, through the conversion process, a single GFF3 file is converted to thousands of piecemeal JSON files, thus putting a heavy burden on the ability to back up and store data.

In the recently released JBrowse version (v1.12.3), support for indexed GFF3 files has been added[4]. In this strategy, the GFF3 file is compressed with bgzip and indexed with tabix[5], which generates only two data files: a compressed file (.gz) and an index file (.tbi). Compared with the traditional processing protocol, the whole compression and index process could be finished within a few seconds even for large datasets such as the human genome annotation data (Additional file 1). The tabix tool requires GFF3 files to be sorted by chromosomes and positions, which could be performed in the GNU sort program or the GenomeTools[6] package (see [7]). When dealing with feature lines in the same chromosome and position, both of the tools would sort them in an ambiguous way that usually results in parent features being placed behind their children (Figure 1A). Although this is still valid in tabix indexing, it would causing erroneous rendering in JBrowse[8] (Figure 1A). Currently there is no additional options or arguments for current tools to break such tied features by parent-child relationship. In the absence of a suitable bug fix to JBrowse, an alternative sorting tool is needed to resolve this problem.

Here, we present GFF3sort, a novel tool to sort GFF3 files for tabix indexing. Compared with GNU sort and GenomeTools, GFF3sort produces sorting results that could be correctly rendered by JBrowse while still keeps enough efficiency. We anticipate that GFF3sort will be a useful tool to help with processing and visualizing genome annotation data.

## Implementation

GFF3sort is a script written in Perl. It uses a hash table to store the input GFF3 annotation data (Figure 1B). For each feature, the chromosome ID and the start position are stored in the primary and secondary key, respectively. Features with the same chromosome and start position are grouped in an array in the same order of their appearance in the original GFF3 data. After sorting the hash table by chromosome IDs and start positions, GFF3sort implemented two modes to sort features within the array: the default mode and the precise mode (Figure 1B). In most situations, the original GFF3 annotations produced by genome annotation projects have already placed parent features before their children. Therefore, GFF3sort returns the feature lines in their original order, which is the default behavior. In some situations where orders in the input file has not obeyed the parent-child relationship, GFF3sort would sort them according to the parent-child topology using the sorting algorithm of directed acyclic graph[9], which is the most precise behavior but costs a little more computational source.

In order to test the performance of GFF3sort, the GFF3 annotation files of seven species, *Saccharomyces cerevisiae* (R64 −1 −1), *Aspergillus nidulans* (ASM1142v1), *Chlamydomonas reinhardtii* (INSDC v3.1), *Drosophila melanogaster* (BDGP6), *Arabidopsis thaliana* (Araport11), *Rattus norvegicus* (Rnor_6.0), and *Homo sapiens* (GRCh38), were downloaded from the ENSEMBL database [10]. All the tests were conducted on a SuperMicro® server equipped with 80 Intel® Xeon® CPUs (2.40GHz), 128 GB RAM, and running the CentOS 6.9 system. By default, CentOS 6.9 carries GNU sort v8.4, a relatively old version released in 2010. Therefore, we downloaded and installed a new version (v8.28) from the official repository of GNU Coreutils[11]. Both the old and the new version of GNU sort would be used in performance test.

## Results and Discussion

GFF3sort takes a GFF3 file as its input data and returns a sorted GFF3 file as output. Several optional parameters are provided such as turning on the precise mode, sorting chromosomes in different ways and properly dealing with inline FASTA sequences. Element models sorted by GFF3sort can be correctly rendered by JBrowse (Figure 1C).

Besides the fixation of JBrowse rendering, GFF3sort has also other advantages over traditional tools. Compared with the GNU sort program, GFF3sort can properly deal with GFF3-specific lines or directives that are preceded by the '##' symbol, such as the topmost GFF version line and the heading sequence-region line. Compared with the GenomeTools, GFF3sort runs significantly faster (Additional file 1). In the default mode, GFF3sort saves ∼70% running time in our seven test datasets. The precise mode takes longer time but still runs faster than GenomeTools, especially for large annotation data such as human. While keeping a high running speed, the memory consumption is still acceptable (Additional file 1). For the largest annotation dataset (the GRCh38 annotation version of human) with a ∼400MB GFF3 file, the memory usage of GFF3sort is ∼758MB, ∼40% less than GenomeTools.

## Conclusions

In conclusion, GFF3sort is a novel tool to sort GFF3 files for tabix indexing and therefore can be used to visualize annotation data in JBrowse appropriately. It has a fast running speed compared with similar, existing tools. We anticipate that GFF3sort will be a useful tool to simplify data processing and visualization.

**Figure 1.**
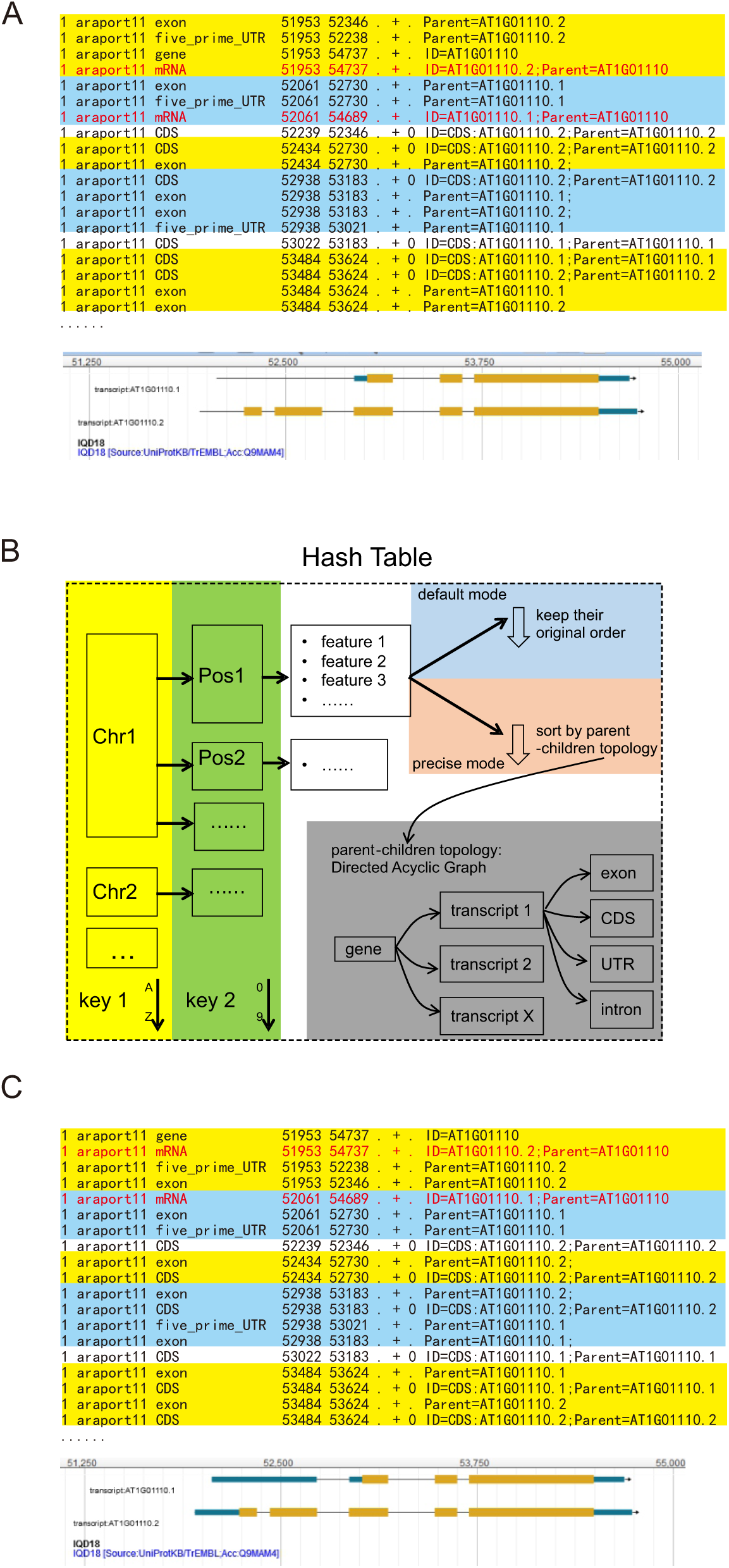
The motivation for, outlines of, and action effects of GFF3sort. A) An example of incorrectly sorted GFF3 data and its snapshots in JBrowse. Blocks with the same start position are marked in blue-yellow stripes. The two lines (mRNA) marked in red were placed after their sub-features (exon or UTR). Such incorrect placement leads to losing the first exon in JBrowse rendering results. See Additional file 2 for the full annotation lines. B) Outlines of GFF3sort. C) An example of correctly sorted data by GFF3sort and its snapshots in JBrowse. In this example, the two lines (mRNA) marked in red were correctly placed before their sub-features, allowing JBrowse to render them properly.

## Additional files

Additional file 1: Benchmark data. This file displays: 1) the detailed running time of GFF3 -to -JSON conversion and the bgzip-tabix process on our test datasets; 2) the detailed running time and 3) memory usage of GFF3sort, GNU sort (v8.4 and v8.28), and GenomeTools on our test datasets. (PDF)

Additional file 2: The full GFF3 annotation lines used in Figure 1A and C. It is the gene AT1G01110 extracted from the *Arabidopsis thaliana* (Araport11) annotation files. It includes three plain-text files: `raw.gff3, GNUsort.gff3` (Figure 1A)and `GFF3sort.gff3` (Figure 1C). (ZIP)

## List of abbreviations

JBrowse: JavaScript-based genome browser

GFF3: General Feature Format, version 3

JSON: JavaScript Object Notation

HTML5: HyperText Markup Language, version 5

## Declarations

### Ethics approval and consent to participate

Not applicable.

### Consent for publication

Not applicable.

### Availability of data and material

Project name: GFF3sort

Project home page: <u>https://github.com/billzt/gff3sort</u>

Operating system(s): Linux

Programming language: Perl

Other requirements: No

License: No restrictions for academic users.

Any restrictions to use by non-academics: license needed

### Competing interests

The authors declare that they have no competing interests.

### Funding

This work is supported by grants from the National Science and Foundation of China (Grant No. 31771850) and the Ministry of Agriculture of China (Grant No. 2016ZX08005004).

### Authors’ contributions

SG, RZ, and TZ initiated the idea of the tool and conceived the project. TZ designed the tool and analyzed the data. CL and ZM helped to test the tool. TZ wrote the paper. All authors read and approved the final manuscript.

## Acknowledgements

We thank Dr. Miklos Csuros and other anonymous reviewers for their helpful comments.

